# Tuning starch granule size distributions in durum wheat using genetic variation at a single locus

**DOI:** 10.1101/2025.03.10.642400

**Authors:** Brendan Fahy, Jiawen Chen, David Seung

## Abstract

The size distribution of starch granules in wheat grains influences bread- and pasta-making quality, as well as nutritional properties. Here, we demonstrate that in durum wheat, wide variation in starch granule size distributions can be induced through missense mutations at a single genetic locus encoding the MYOSIN RESEMBLING CHLOROPLAST PROTEIN on chromosome 6A (*TtMRC-A1*). We isolated 29 independent TILLING mutants in durum cultivar Kronos, each harbouring a different missense mutation that causes an amino acid substitution in the MRC protein. Compared to the B-type granule content of wild-type Kronos (24%), six of the missense lines had significant increases in B-type granule content (33-42%), although not to the extent observed in the *mrc-1* mutant (58%) which carries a premature stop codon mutation. Notably, one missense line had significantly decreased B-type granule content (15%), demonstrating that mutations in *TtMRC-A1* can achieve both increases and decreases in B-type granule content. In these lines, A-type granule size decreased as B-type granule content increased, and Rapid Visco Analysis on selected lines demonstrated that both B-type granule content and A-type granule size strongly correlated with pasting parameters (e.g., peak viscosity and pasting temperature). However, strong correlations between pasting properties and A-type granule size were still observed after removing most of the B-type granules via sieving, indicating that A-type granule size is the primary contributor to the observed variation in pasting properties. Overall, we demonstrate that mutations at *TtMRC-A1* can greatly extend the range of granule size distributions in durum wheat, creating useful alterations in starch properties.

**Key Message:** Different missense mutations in *TtMRC-A1* can be used to fine tune granule size distributions in durum wheat grains, creating useful alterations in starch properties.

## Introduction

Starch granule size distributions influence both the functional and nutritional properties of starch (Li et al. 2021; Lindeboom et al. 2004; Qi and Tester 2016). In grains of wheat and other Triticeae crops, there is a bimodal distribution of starch granule size, consisting of large A-type and small B-type granules. A-type granules are lenticular in shape and are initiated during the early phases of grain development, whereas B-type granules are spherical and initiated about 10-15 days after the A-type granules (Buttrose 1960; Evers 1971).

The effect of starch granule size distributions on end-use quality has been studied using the variation in this trait among different wheat cultivars, or by creating more extreme variation through flour reconstitution after granule size fractionation. For breadmaking, there was an optimum amount of B-type granules (often referred to as B-type granule content) for crumb grain and texture, which depended on the protein content of the grain (Lelievre et al. 1987; Park et al. 2005; Park et al. 2009). This suggests that starch granule size distributions affect the protein-starch interactions that are important for rheological properties and bread quality. In pasta-making experiments with reconstituted flours, increased B-type granule content enhanced pasta firmness (Soh et al. 2006). In addition to textural qualities, starch granule size affects nutritional properties. In general, larger starch granules are digested more slowly in vitro than smaller granules due to their reduced surface area per volume ratio (Dhital et al. 2010). Indeed, native A-type granules are digested more slowly in vitro than B-type granules (Colussi et al. 2021; Moon and Kweon 2024). The extent to which these trends translate to cooked starch is not known, but it is important to note that granules do not completely gelatinise when cooked in a food matrix. For example, cooked pasta contains both intact and incompletely gelatinised starch granules, particularly towards the internal regions of the noodle (Zou et al. 2015). Thus, increasing the amount of A-type granules in durum wheat, as well as their size, could boost the amount of resistant starch in pasta.

Genetics offers a means to modify granule size distributions in wheat. The range of B-type granule content in durum wheat was successfully increased through crossing with *Triticum dicoccoides* lines that have substantial variation in this trait (Sissons and Egan 2023). However, for the range of B-type granule content achieved (22-44%), there were no relationships between B-type granule content and pasta quality parameters, suggesting that a wider range of variation could be necessary for achieving the positive effects seen in flour reconstitution experiments. We reasoned that the recent identification of genes required for B-type granule synthesis may help achieve this wider range. For example, reduced activity of *B GRANULE CONTENT1 (BGC1)*, or knockout of ⍺-glucan *PHOSPHORYLASE (PHS1)*, can lead to wheat with reduced B-type granule content (Chia et al. 2020; Kamble et al. 2023; Saccomanno et al. 2022).

Another promising gene for altering B-type granule content encodes the MYOSIN RESEMBLING CHLOROPLAST PROTEIN (MRC). MRC (also referred to as PROTEIN INVOLVED IN STARCH INITIATION, PII1) is a long coiled-coil protein which promotes starch granule initiation in Arabidopsis leaves (Seung et al. 2018; Vandromme et al. 2019; Vandromme et al. 2023). In wheat, MRC represses B-type granule initiation during early grain development (Chen et al. 2024). Premature stop codon mutations in MRC in the durum wheat Kronos dramatically increased the B-type granule content, up to 68%. This increase resulted from the early initiation of B-type granules during grain development, which also led to a concomitant decrease in the average size of A-type granules. MRC is a particularly useful target in durum wheat because it has only one functional homeolog (*TtMRC-A1* on chromosome 6A) due to the chromosome 6B homoeolog becoming a pseudogene, allowing manipulation of B-granule content through mutations at a single locus. Here, we screened B-type granule content in 29 durum wheat lines, each carrying different missense mutations in the MRC locus. We identified lines with both increased and decreased B-type granule content, generating a remarkable range of B-granule content (15-58%) within the single isogenic background of cultivar Kronos. We demonstrate using the Rapid Visco Analyser (RVA) that the lines at the extremes of this variation have substantial changes in starch pasting properties. Overall, we demonstrate that mutations at a single genetic locus can be used to tune granule size distributions in durum wheat, offering ideal material for testing how granule size affects functional and nutritional quality.

## Materials and Methods

### Plant materials and growth

Seeds of wheat TILLING lines were obtained from the John Innes Centre (JIC) Germplasm Resources Unit. Mutants from Chen et al. (2024) include *mrc-1* (referred to as the BC2 *aabb* line in the original publication) and *mrc-2*. KASP markers used to genotype the mutations are provided in Supplemental Table 1. For experiments 1 and 2, plants were grown in a Controlled Environment Room fitted with fluorescent lamps supplemented with LED panels. The chamber was set to provide 16 h light (400 µmol photons m_−2_ s_−1_) at 20°C and 8 h dark at 16°C, with constant relative humidity (65%). For bulking grains for the RVA experiment, 96 plants per genotype were grown in 2 L pots in the glasshouse (4 plants per pot) until maturity and seed harvest. The glasshouse was set to provide a minimum 16 h of light at 20°C and 16°C during the dark. All plants were grown in John Innes cereal mix - 65% peat, 25% loam, 10% grit, 3 kg/m_3_ dolomitic limestone, 1.3 kg/m_3_ pg mix, and 3 kg/m_3_ osmocote exact.

### Small-scale starch extraction

Small scale starch extraction was used for the analyses in Experiments 1 and 2. Briefly, seeds (3 seeds per replicate) were cracked with a plier and soaked overnight in 0.5 M NaCl (0.75 mL) at 4°C, then homogenised for 6 min in a MM-300 ball mill (Retsch)(set to 25 Hz) with a 5 mm steel ball. The homogenate was filtered through Miracloth, then through a pluriStrainer Mini with 70 µm nylon mesh (Pluriselect). Starch in the filtrate was pelleted at 18,000*g*, then resuspended in 300 µL ddH_2_O and placed over a 1 mL cushion of 90% (v/v) buffered Percoll solution (prepared in 50 mM Tris-HCl, pH 8). The starch was spun through the Percoll at 2500*g* for 15 min. The starch pellet was then washed 3 times in 1 mL of 2% SDS, then once in 1 mL acetone, spinning at 18,000*g* for 5 min after each resuspension. The starch was air dried.

### Large-scale starch extraction and fractionation

Large-scale starch extraction from bulked grains was used to produce starch for the RVA analyses. Grains (20 g per extraction) were soaked overnight in ddH_2_O at 4°C, before homogenising in a mortar and pestle. The homogenate was then washed with excess water through muslin, followed by Miracloth. The starch in the filtrate was collected by centrifugation at 3,000*g* for 5 min, then resuspended in 8 mL ddH_2_O before placing over a 36 mL cushion of 90% (v/v) buffered Percoll solution (prepared in 50 mM Tris-HCl, pH 8). The starch was spun through the Percoll at 2500*g* for 15 min. The starch pellet was washed 3 times in 20 mL of 2% SDS, then once in 20 mL acetone, spinning at 3,000*g* for 5 min after each resuspension. The starch was air dried.

To remove B-type granules from the starch preparation, purified starch was suspended in 70% ethanol, and then filtered through a pluriStrainer fitted with a 10 µm nylon mesh (Pluriselect). Filtration was assisted by placing the strainers over a magnetic stirrer, and adding a stir bar within each strainer above the nylon mesh to prevent starch from settling and blocking the filters. Once filtration had completed, more 70% ethanol was added to the strainers to wash remaining B-type granules through the nylon mesh. After filtration and washing was completed, starch enriched in A-type granules in the retentate was collected by placing the strainers into a bottle with 70% ethanol and shaking to resuspend the starch. The starch was then collected by pelleting at 3,000*g* for 5 min, washed once in acetone, then air dried.

### Total starch quantification and grain morphometrics

For total starch quantification, grains (5 grains per replicate) were crushed with pliers into a 2 mL centrifuge tube, then milled into flour using a MM-300 ball mill (Retsch)(set to 30 Hz) with a 5 mm steel ball. Quantification was carried out using the Total Starch HK Kit (Megazyme) according to the manufacturer’s instructions, except the starting material was adjusted to 5-10 mg of flour, and all assay volumes were reduced by 6X. Grain size was quantified using the MARViN seed analyser (Marvitech GmbH, Wittenburg).

### Scanning Electron Microscopy (SEM)

Purified starch suspended in water was air-dried onto a glass coverslip attached onto an SEM stub. Stubs were sputter coated with gold and observed using a Gemini 300 FEG SEM microscope (Zeiss).

### Coulter counter analysis

Purified starch granules were resuspended in Isoton II electrolyte (Beckman Coulter) and analysed on a Multisizer 4e Coulter counter (Beckman Coulter), fitted with a 70 µm aperture and 100 mL beaker. We measured a minimum of 50,000 particles per run. Granule size parameters were derived using curve fitting scripts, described in McNelly et al. (2024)

### Rapid Visco Analysis

Rapid Visco Analysis (RVA) was carried out on an RVA Tecmaster instrument (Perten, Waltham, MA, USA) running the preinstalled general pasting method, according to AACC Method 76-21. Analyses were performed with 2 g purified starch in 25 mL of water.

## Results

### Variation in granule size distributions in durum wheat through missense mutations in MRC

Using the wheat TILLING mutant database on Ensembl plants (Krasileva et al. 2017), we identified a total of 45 different mutant lines in the tetraploid durum wheat Kronos that carried different missense mutations in *TtMRC-A1* (Table 1). The amino acid substitutions in these lines were in different positions along the length of the MRC protein, and were predicted to range from tolerated to deleterious using both the SIFT and PROVEAN programs (Choi and Chan 2015; Ng and Henikoff 2003). Seeds from all 45 mutant lines (either homozygous or heterozygous for the mutations) were obtained from the Germplasm Resource Unit of the John Innes Centre, and KASP genotyping markers were designed for each of the mutations. For our experiments, we successfully genotyped and isolated 29 homozygous lines (Table 1). The remaining 16 lines were excluded, either because the mutation could not be successfully genotyped using KASP markers, due to poor growth of the mutant line (indicating the presence of background mutations), or because not enough homozygous seeds were obtained in time for the first experiment.

**TABLE 1.** Kronos *mrc* mutants used in this study.

| **TABLE 1** |  |  |  |  |
| --- | --- | --- | --- | --- |
| Kronos *mrc* mutants used in this study |  |  |  |  |
| Genotype | Mutation | SIFT score | PROVEAN score | Comments: |
| **Missense mutants included in the study (29)** |  |  |  |  |
| K3745 | A19T | 0.18 | -0.856 | Exp. 1 |
| K3543 | G63S | 0.7 | 0.926 | Exp. 1 |
| K1435 | S92N | 0.06 | 0.004 | Exp. 1 |
| K2447 | L149F | 0* | -3.944* | Exp. 1 |
| K2937 | A178T | 0.33 | -2.114 | Exp. 1; Exp. 2 |
| K2124 | R196K | 0.75 | 0.12 | Exp. 1 |
| K2205 | E213K | 0.15 | -2.437 | Exp. 1 |
| K3499 | R230C | 0.16 | -2.755* | Exp. 1 |
| K1431 | E225K | 0* | -3.911* | Exp. 1 |
| K2397 | R230C | 0.04* | -2.755* | Exp. 1 |
| K4060 | R255K | 0.38 | 0.05 | Exp. 1; Exp. 2 |
| K0598 | L289F | 0* | -3.961* | Exp. 1; Exp. 2 |
| K3533 | A346V | 0.13 | -3.278* | Exp. 1 |
| K2981 | A356T | 0.11 | -0.575 | Exp. 1 |
| K0775 | L394F | 0* | -3.5* | Exp. 1 |
| K0265 | M412I | 0.03* | -2.153 | Exp. 1; Exp. 2 |
| K3424 | E491K | 0.01* | -3.267* | Exp. 1 |
| K2408 | S503N | 1 | 3.094 | Exp. 1; Exp. 2 |
| K1232 | S539N | 1 | 1.528 | Exp. 1 |
| K3889 | D581N | 0.13 | -2.011 | Exp. 1; Exp. 2 |
| K3505 | E606K | 0.22 | -1.96 | Exp. 1; Exp. 2 |
| K2485 | A625T | 0.01* | -2.79* | Exp. 1; Exp. 2 |
| K4297 | P656L | 0.13 | -1.238 | Exp. 1 |
| K3416 | S667N | 0.24 | 0.188 | Exp. 1 |
| K2348 | E679K | 0.02* | -0.233 | Exp. 1 |
| K2096 | P681S | 0.01* | 0.835 | Exp. 1 |
| K2821 | A687V | 0.13 | -3.182* | Exp. 1 |
| K3666 | D708N | 0.02* | -1.631 | Exp. 1 |
| K4193 | S726N | 0.46 | -0.47 | Exp. 1 |
| **Missense mutants excluded in the study (16)** |  |  |  |  |
| K4533 | A79V | 0.01* | -3.263* | Insufficient seeds |
| K2450 | A155T | 0.12 | -0.42 | Poor growth |
| K4618 | R196K | 0.75 | 0.12 | Insufficient seeds |
| K0565 | S329F | 0.01* | -3.214* | Unable to genotype |
| K0665 | A356T | 0.11 | -0.575 | Insufficient seeds |
| K3286 | S375F | 0.01* | -5.338* | Poor growth |
| K2976 | L409F | 0.07 | -0.445 | Insufficient seeds |
| K1112 | E413K | 0.15 | -3.465* | Poor growth |
| K3145 | E425K | 0.04* | -3.8* | Poor growth |
| K3657 | T500I | 0.07 | -1.68 | Unable to genotype |
| K1161 | T525I | 0.1 | -3.878* | Poor growth |
| K3661 | D575N | 0.86 | 0.287 | Poor growth |
| K4205 | L591F | 0.24 | -0.127 | Poor growth |
| K3137 | A603T | 1 | 2.048 | Unable to genotype |
| K0338 | R700K | 0.48 | -0.325 | Insufficient seeds |
| K4082 | D743N | 0.52 | -1.675 | Unable to genotype |
| **Premature stop codon mutants included in the study (2)** |  |  |  |  |
| *mrc-1* | W258STOP | - |  | Exp. 1; Exp. 2 |
| *mrc-2* | W550STOP | - |  | Exp. 1 |
| SIFT and PROVEAN scores that are considered damaging (using default cut-off values of 0.05 and -2.5 for each program, respectively), are marked with an asterisk (*). For lines included in the study, the experiment(s) in which phenotypic analyses were conducted is provided in the comments. For lines excluded, the reason for exclusion is provided in the comments. |  |  |  |  |

In our first experiment (Experiment 1), we screened the 29 homozygous mutant lines for alterations in starch granule size distribution in the endosperm. Starch was extracted from mature grains harvested from n = 3-4 individual plants per line, which were all grown together in a single growth experiment in the glasshouse. For comparison, the *mrc-1* and *mrc-2* mutants which carry premature stop codons in *TtMRC-A1* were also included (Chen et al. 2024)(Table 1). Granule size distributions were analysed with the Coulter counter, and subsequent curve fitting was used to derive B-type granule content (McNelly et al. 2024). The *mrc-1* mutant had the highest B-type granule content, whereas *mrc-2* had a relatively moderate increase in B-type granule content compared to *mrc-1*. These results were consistent with Chen et al. (2024) and likely due to the premature stop codon occurring later in the coding sequencing in *mrc-2* compared to *mrc-1* (Table 1). Interestingly, three of the lines with missense mutations had B-type granule content that was in between that of *mrc-1* and *mrc-2*. This included K0598 (referred to as *mrc-3* in Chen et al. 2024), K4060 and K3889. Strikingly, some of the lines had a lower B-type granule content than the wild type, with the lowest (K2485) having a value almost half that of the wild type.

Although we observed substantial variation in B-granule content in this experiment, a one-way ANOVA with Tukey’s HSD test showed that only the B-type granule content values of *mrc-1* and K0598 were significantly different from the WT. This is due to low statistical power within the experiment, caused by the large number of lines being compared within the experiment, and the relatively low numbers of replicates per line. However, under pairwise two-tailed t-tests against the wild type, six of the missense lines (K3505, K2937, K2408, K4060, K3889, K0598), alongside *mrc-1* and *mrc-2*, had significantly higher B-type granule content than the wild type, while two lines (K0265, K2485) had significantly lower B-type granule content. Experiment 1 was therefore successful as a screen that provided candidate lines with altered B-type granule content, but the lack of statistical power meant that more replicates were required for further characterisation.

Therefore, in Experiment 2, we selected the six candidate lines that had higher B-type granule content than the wild type under pairwise t-tests, along with both lines with lower B-type granule content, and grew them for a second generation in the glasshouse with a larger number of replicates (n = 7-8 plants per line). Mature grains were harvested for starch granule analyses. The smaller number of genotypes and more replication compared to Experiment 1 increased statistical power, as one-way ANOVA and Tukey’s HSD post-hoc test revealed more lines with significantly altered B-type granule content than the previous experiment (Figure 2A). Like Experiment 1, *mrc-1* and K0598 had the largest B-type granule content, at 58% and 42% respectively. The value obtained for *mrc-1* was slightly lower than the 67% increase we reported in Chen et al. (2024). Four of the missense mutants (K4060, K3889, K2408 and K2937) had an intermediate increase in B-type granule content (33-36%), that was significantly different from the wild type (24%). Also similar to Experiment 1, K2486 and K0265 were the only lines to have lower B-type granule content, but the effect was significant only for K2485 (15%). Overall, in this experiment, the different *mrc* mutations achieved a wide range of B-type granule content between 15-58%.

**Figure 1:**
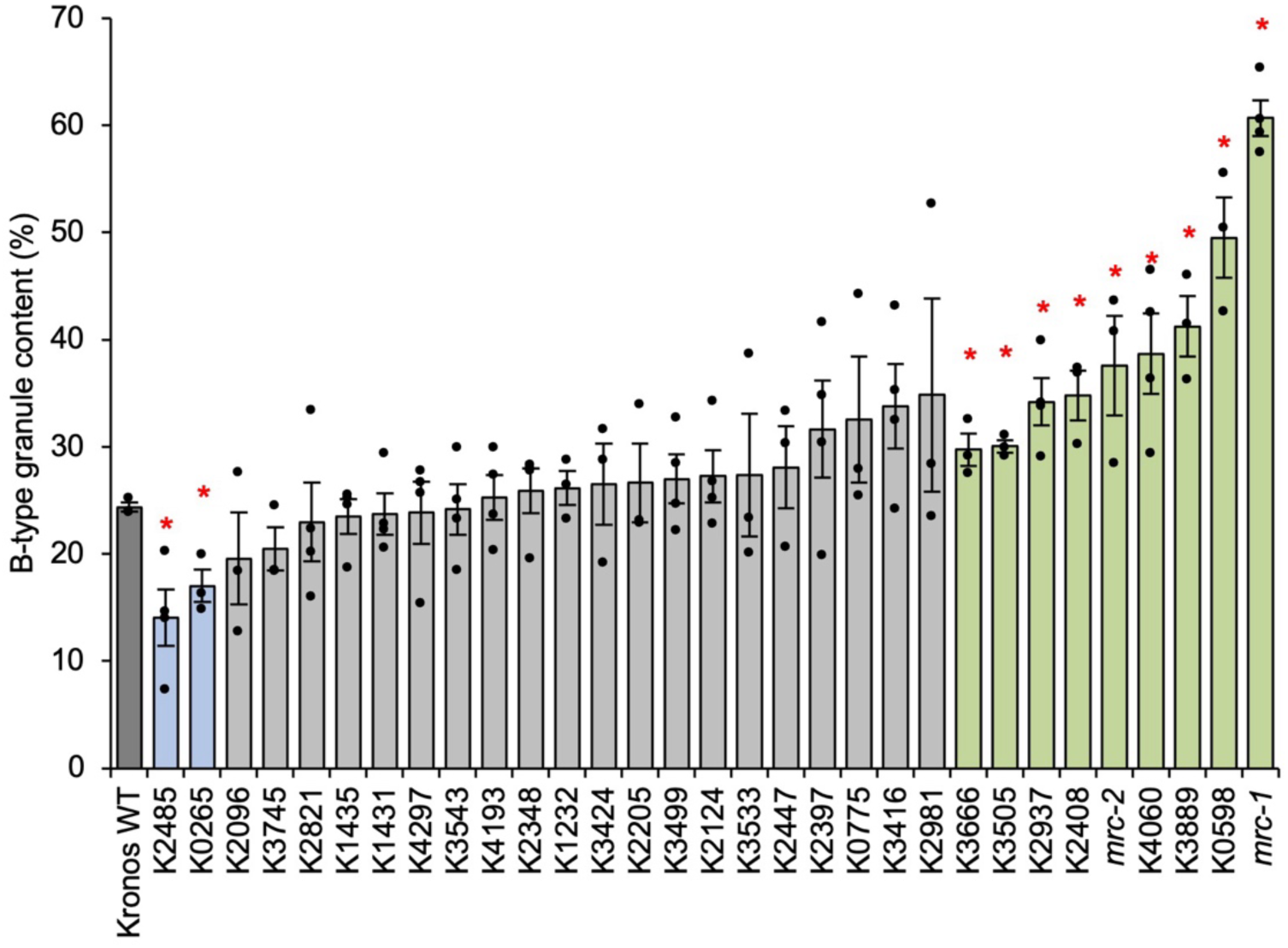
B-granule content from initial screening of Kronos TILLING mutants (Experiment 1). Starch was extracted from mature grains and B-type granule volume was calculated using Coulter counter and curve-fitting analyses. Values are the mean ± standard error of the mean (SEM) from *n*=3-4 biological replicates, where each replicate (shown as an individual data point) was prepared from grains harvested from a separate plant. Values marked with an asterisk are significantly different to the Kronos wild type (WT) under a pairwise two-tailed t-test (p < 0.05).

**Figure 2:**
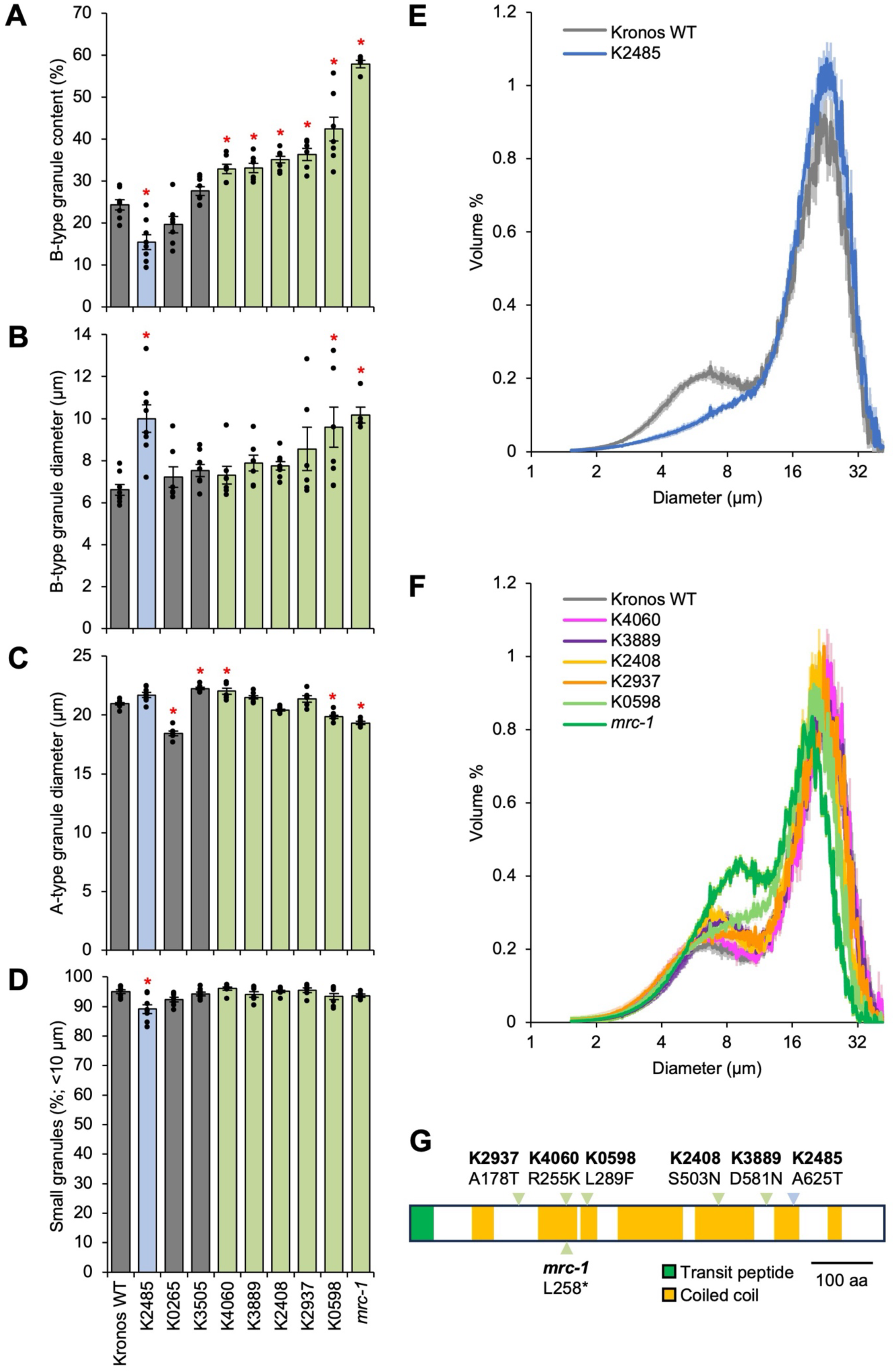
Granule size parameters from Kronos TILLING mutants in Experiment 2. Coulter Counter analyses were carried out on starches extracted from mature grains, and granule size parameters were calculated using curve-fitting analyses. **(A)** B-type granule content (by relative volume). **(B)** Mean B-type granule diameter. **(C)** Mean A-type granule diameter. **(D)** Relative number of small granules, defined as granules < 10 µm. For panels A-D, values are the mean ± standard error of the mean (SEM) from *n*=7-8 biological replicates, where each replicate (shown as an individual data point) was prepared from grains harvested from a separate plant. Values marked with an asterisk are significantly different to the Kronos wild type (WT) under a one-way ANOVA and Tukey’s HSD test (p < 0.05). **(E)** and **(F)** Volumetric granule size distribution from the Coulter counter for the K2485 line with low B-type granule content (panel E), and for lines with high B-type granule content (panel F). These are from the same analyses as those presented in A-D. The solid line represents the mean distribution from *n*=7-8 biological replicates, and the shading represents the SEM. Data were acquired with logarithmic bin spacing and are shown on a logarithmic x-axis. **(G)** Schematic representation of the *Tt*MRC-A1 polypeptide, with locations of amino acid substitutions in the different mutants indicated. Locations of mutations that significantly increased B-type granule content are marked with a green arrowhead, while the location of the K2485 mutation that significantly decreased B-type granule content is marked with a blue arrowhead.

We then assessed the mean size of the A- and B-type granules, another granule size parameter from the Coulter counter and curve-fitting analysis. Consistent with Chen et al. (2024), the *mrc-1* knockout mutant and K0598 had significantly larger B-type granules than the wild type (Figure 2B), and significantly smaller A-type granules (Figure 2C). We previously proposed that the smaller A-type granules in these mutants is due to the increased amount of B-type granules competing for substrates with A-type granules (Chen et al. 2024). However, the four lines with intermediate increases in B-type granule content (K4060, K3889, K2408 and K2937) did not show consistent decreases in A-type granule size, suggesting that such moderate increases are not sufficient to cause detectable decreases in A-type granule size.

Interestingly, the mean size of B-type granules was significantly larger in line K2485 compared to the wild type, despite its significant reduction in total B-type granule content (Figure 2B), which is only possible if this line has fewer B-type granules by number. We therefore calculated the relative number of small granules from the Coulter counter data, defining small granules as < 10 µm, which would mostly contain B-type granules. K2485 was the only line with a significant change in this parameter, indeed having significantly fewer small granules (Figure 2D). K2485 therefore makes fewer but larger B-type granules compared to the wild type, and an overall decrease in B-type granule content by volume. This was reflected in the drastically altered granule size distribution curve compared to the wild type, where the B-type granule peak was much less prominent in the mutant (Figure 2E). This starkly contrasted all other *mrc* mutants, which had more prominent B-type granule peaks (Figure 2F).

To complement the analysis of granule morphology using the Coulter counter, we visually examined the starch granules in these lines using Scanning Electron Microscopy. Although quantitative differences in granule size distribution are difficult to discern using microscopy, we observed that all lines had typical lenticular A-type and round B-type granule morphology (Supplemental Figure 1). In addition, we measured total starch content in these selected lines. With the exception of one line (K2408) that had a small reduction in starch content compared to the wild type, all other lines had no significant changes in starch content (Supplemental Figure 2A), indicating that the altered granule size distributions were largely independent of starch content. To check for effects on grain size, we measured the Thousand Grain Weight (TGW) using the Marvin seed analyser (Supplemental Figure 2B). For unknown reasons, four of the examined lines had elevated TGW (K3505, K4060, K3889 and K2408), but this did not correlate with the variation in starch granule size, as the lines with the most extreme changes in starch granule size (e.g., K2485, K0598 and *mrc-1*) did not have altered TGW.

Overall, we discovered that mutations in *TtMRC-A1* can be used to both increase and decrease B-type granule content without affecting total starch content or grain size, making it an ideal target gene for achieving a wide range of granule size distributions in durum wheat. The amino acid substitutions that led to these changes followed no obvious pattern. Those that increased B-type granule content were scattered along the length of the *Tt*MRC-A1 polypeptide and were not necessarily in a coiled coil region, while the A625T mutation in K2485 that decreased B-type granule content was in a coiled coil towards the C-terminus (Figure 2G).

### The altered granule size distribution affects starch physicochemical properties in the mutants

Our durum lines presented a unique opportunity to study how variation in granule size parameters affects starch functional properties within a single cultivar. We therefore selected lines for bulking grains and for conducting starch functional analyses. We chose the line with the highest B-type granule content (*mrc-1*), the K0598 line with an intermediate increase in B-type granule content, and the K2485 line with reduced B-type granule content, together with the wild-type Kronos background. Grains were bulked in the glasshouse and we used a large-scale starch extraction method to obtain purified starch. Coulter counter analyses showed that the large-scale starch preparation showed similar trends to what we observed with the small-scale experiments, with K2485 having a lower B-type granule peak than the wild type, whereas both K0598 and *mrc-1* had higher B-type granule peaks than the wild type (Figure 3A). However, the measured B-type granule content for all genotypes was higher than what we observed in the small-scale experiments, ranging from 31% in K2485 to 64% in *mrc-1* (Supplemental Figure 3), although the value for *mrc-1* was still in a range that was expected for this genotype from our previous work (Chen et al. 2024).

**Figure 3:**
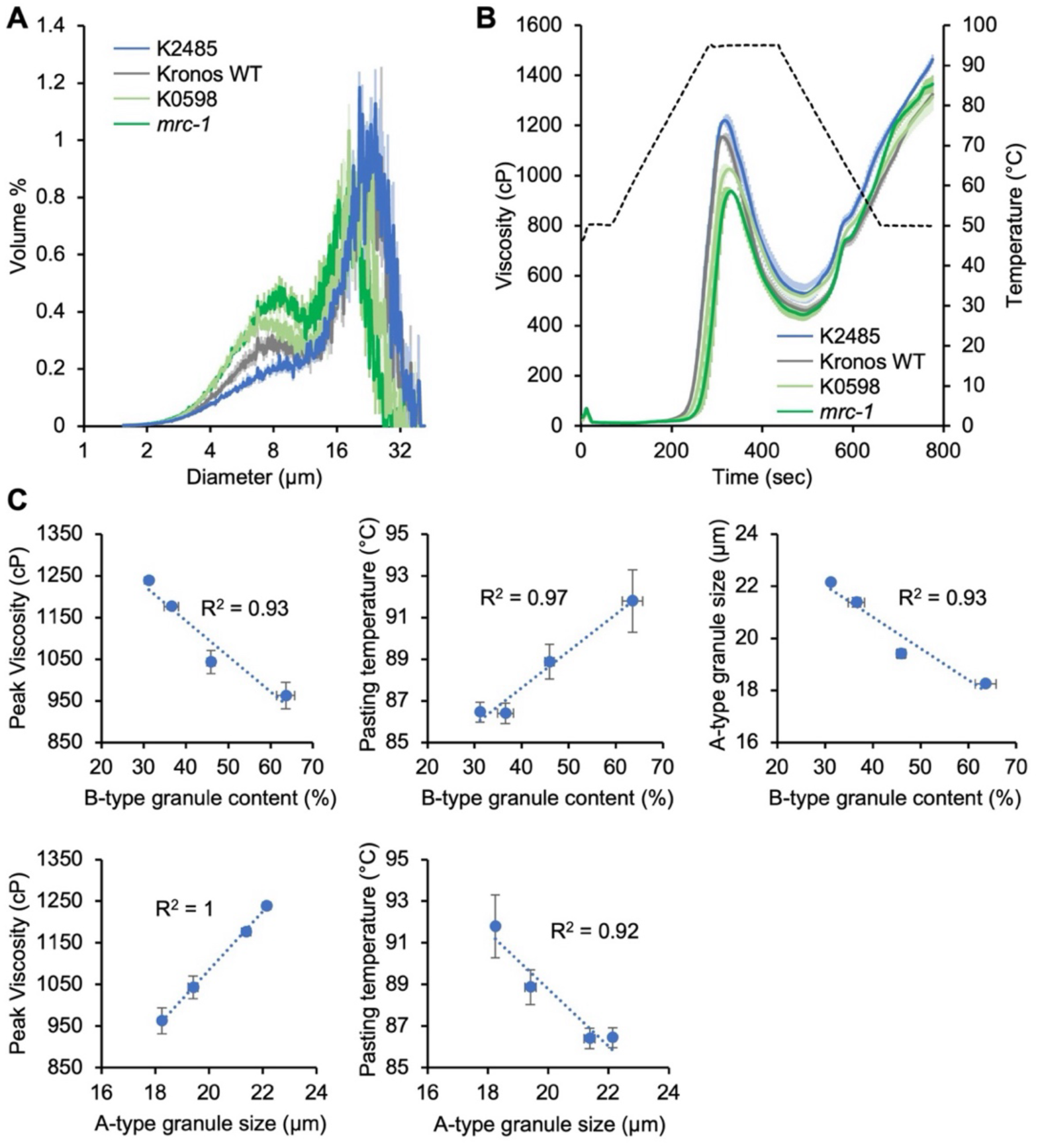
Mutants in *TtMRC-A1* with altered granule size distributions have altered pasting properties. **A)** Granule size distributions of starch samples from bulked grains, measured using a Coulter counter. Data were acquired with logarithmic bin spacing and are shown on a logarithmic x-axis. **B)** Rapid Visco Analysis (RVA) showing pasting profiles of the starches shown in panel A. For both panels A and B, the solid line represents the mean from n = 3 replicate measurements, while the shading indicates the standard error of the mean (SEM). **C)** Correlation plots between granule size and pasting parameters. The R^2^ value is shown on all panels.

We then analysed starch pasting parameters using the Rapid Visco Analyser (Figure 3B; Table 2). The viscograms showed substantial variation between the tested lines: both lines with higher B-type granule content (*mrc-1* and K0598 lines) had lower peak viscosity than the wild type, while the K2485 line with lower B-type granule content had higher viscosity. To identify which pasting (RVA) parameters were most dependent on granule size distributions, we performed pairwise linear regressions between each pasting parameter and each granule size parameter, using R_2_ values to quantify the strength of these relationships (Supplemental Figure 4). The strongest relationships with B-type granule content were a negative correlation with peak viscosity (R_2_ = 0.93) and positive correlation with pasting temperature (R_2_ = 0.97)(Figure 3C). However, as expected from our results in Experiment 2, B-type granule content also strongly, negatively correlated with A-type granule size (R_2_ = 0.93)(Figure 3C); and thus all RVA parameters that negatively correlated with B-type granule content were also positively correlated with A-type granule diameter, and vice versa (Supplemental Figure 4). For example, A-type granule size was strongly positively correlated with peak viscosity (R_2_ = 1) and negatively with pasting temperature (R_2_ = 0.92)(Figure 3C). A-type granule size also correlated with breakdown viscosity and peak time (Supplemental Figure 4). B-type granule size correlated to only few pasting parameters, most notably with setback viscosity. Overall, the altered granule size distributions in the selected mutants affected the pasting properties of the starch, but it was not clear whether this was primarily due to variation in B-type granule content, or A-type granule size.

**TABLE 2.** Starch pasting parameters from RVA experiments.

| TABLE 2 |  |  |  |  |  |  |  |
| --- | --- | --- | --- | --- | --- | --- | --- |
| Starch pasting parameters from RVA experiments |  |  |  |  |  |  |  |
| Sample | Peak Visc (cP) | Trough Visc. (cP) | Breakdown Visc. (cP) | Final Visc. (cP) | Setback Visc. (cP) | Peak Time (min) | Pasting temp. (°C) |
| **Whole starch sample** |  |  |  |  |  |  |  |
| Kronos WT | 1176 ± 6 | 461 ± 18 | 715 ± 23 | 1325 ± 11 | 864 ± 28 | 5.24 ± 0.09 | 86.40 ± 0.49 |
| K2485 | 1239 ± 8 | 529 ± 40 | 710 ± 33 | 1464 ± 19 | 936 ± 59 | 5.27 ± 0.10 | 86.45 ± 0.48 |
| K0598 | 1043 ± 27* | 517 ± 22 | 526 ± 49 | 1314 ± 55 | 796 ± 77 | 5.38 ± 0.12 | 88.87 ± 0.83 |
| *mrc-1* | 962 ± 31* | 444 ± 25 | 518 ± 56* | 1366 ± 35 | 922 ± 56 | 5.44 ± 0.16 | 91.78 ± 1.50* |
| **B-type granule-depleted sample** |  |  |  |  |  |  |  |
| Kronos WT | 1202 ± 4 | 627 ± 2 | 575 ± 3 | 1296 ± 10 | 669 ± 8 | 5.49 ± 0.02 | 85.82 ± 0.29 |
| K2485 | 1289 ± 12* | 648 ± 6 | 641 ± 7* | 1453 ± 20* | 805 ± 14* | 5.4 ± 0* | 85.03 ± 0.28 |
| K0598 | 1114 ± 10* | 680 ± 9* | 434 ± 3* | 1267 ± 9 | 587 ± 3* | 5.6 ± 0* | 86.73 ± 0.28 |
| *mrc-1* | 1069 ± 7* | 663 ± 3* | 406 ± 5* | 1313 ± 5 | 650 ± 2 | 5.8 ± 0* | 90.43 ± 0.03* |
| RVA parameters were calculated from the viscograms shown in Figures 3B and 4B. Values are the mean ± standard error of the mean (SEM) from n = 3 replicate measurements. Values marked with an asterisk are significantly different to the Kronos wild type (WT) under a one-way ANOVA and Tukey's HSD test (p < 0.05). |  |  |  |  |  |  |  |

To resolve whether the altered pasting properties were due to the variation in B-type granule content or A-type granule size, we measured pasting parameters on starch samples from the selected mutants after removing most of the B-type granules by sieving the starch through a 10 µm mesh. The retentate after sieving contained starch that was highly enriched in A-type granules, as determined using Coulter counter analysis (Figure 4A). Compared with the original RVA analyses done on the whole starch preparation (Figure 3B), the removal of B-type granules slightly increased peak viscosity in all genotypes, while final viscosity remained similar (Figure 4B; Table 2). Importantly, the relative differences between the genotypes remained similar, with K2485 having higher peak viscosity than the wild type, K0598 having lower peak viscosity than the wild type, and *mrc-1* having the greatest reduction in peak viscosity. The strong correlation between A-type granule size and peak viscosity was retained after sieving, and the correlation between A-type granule size and pasting temperature was also retained, although the R_2_ value was slightly weaker than before sieving (Figure 4C; Supplemental Figure 4). These data suggest that A-type granule size is the primary contributor to the variation in pasting properties in these mutants.

**Figure 4:**
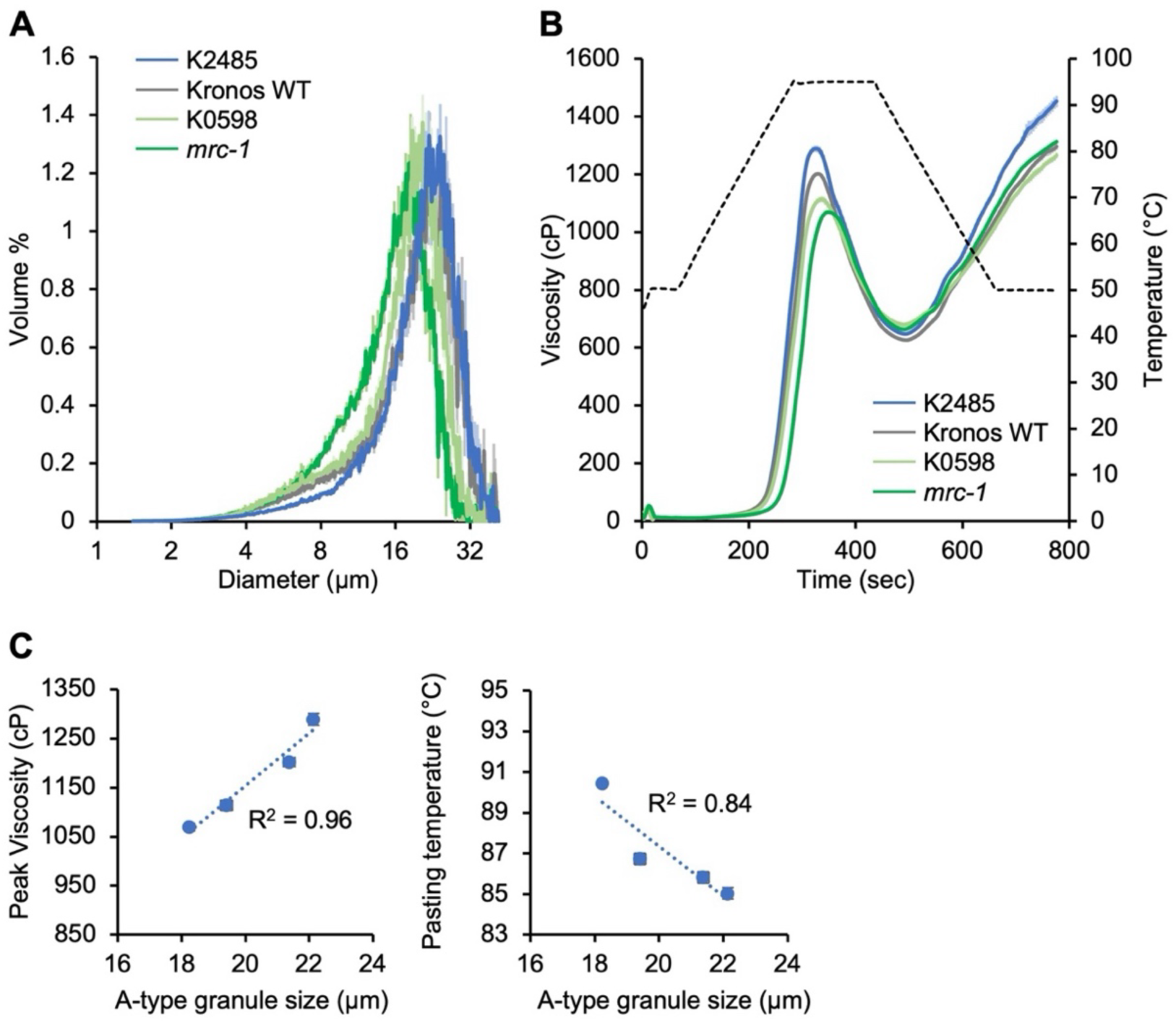
The mutants in *TtMRC-A1* retain altered pasting properties after removal of most B-type granules. **A)** Granule size distributions of starch samples after depletion of B-type granules using a 10 µm filter. Size distributions were measured using a Coulter counter. Data were acquired with logarithmic bin spacing and are shown on a logarithmic x-axis. **B)** Rapid Visco Analysis showing pasting profiles of the starches with depleted B-type granules shown in A. For both panels, the solid line represents the mean from n = 3 replicate measurements, while the shading indicates the standard error of the mean (SEM). **C)** Correlation plots between A-type granule size and pasting parameters. The R^2^ value is shown on both panels.

## Discussion

### Tuning granule size distributions using mutations at a single genetic locus

Here, we demonstrate that starch granule size distributions in durum wheat grains can be tuned at a single genetic locus, *TtMRC-A1*. Mutations at this locus can be used to achieve a wide range of different B-type granule content and A-type granule size. We had previously discovered that *Tt*MRC acts as a repressor of B-type granule initiation during early grain development, such that mutations in *TtMRC-A1* substantially increase B-type granule content (Chen et al. 2024). By screening durum wheat lines carrying *TtMRC-A1* missense mutant alleles, we characterised the effect of random amino acid substitutions along the length of the protein (Table 1). From the 29 missense mutants examined, we identified 5 lines with varying degrees of increased B-type granule content, and unexpectedly one line (K2485) with significantly decreased B-type granule content (Figure 2). The previously characterised *mrc-1* mutant carrying an early premature stop codon (Chen et al. 2024) had the strongest increase in B-type granule content. Using mutations in *TtMRC-A1*, we achieved a range of B-type granule content between 15-58% - where K2485 had the lowest B-type granule content, and *mrc-1* had the highest (Figure 2). This range is greater than what was achieved by crossing in natural variation from *Triticum dicoccoides* (22-44%)(Sissons and Egan 2023).

Our results also demonstrate that altering B-type granule content through *TtMRC-A1* indirectly modulates A-type granule size. We previously proposed that in the *mrc-1* mutant, the B-type granules initiate too early during grain development and hinder A-type granule growth by competing for substrates, thus resulting in smaller A-type granules. Consistent with this, when comparing the mutants with the strongest alterations in B-type granule content (K2495, K0598 and *mrc-1*), we observed a strong negative correlation between B-type granule content and A-type granule size (Figure 3). However, in the lines with intermediate increases in B-type granule content (K4060, K3889, K2408 and K2937), A-type granule content was either not significantly changed, or significantly increased in the case of K4060 (Figure 2). It is likely that such increases in B-type granule content are not sufficient to cause detectable changes in A-type granule size. The reason for the increase in A-type granule size in K4060 is not known but is unlikely to be related to the B-type granules. Interestingly, significant increases in B-type granule size were observed in the two lines with the strongest increases in B-type granule content (K0598 and *mrc-1*), as well as in the K2485 line that had lower B-type granule content. Although this might seem counter-intuitive, the increases in B-type granule size likely occur for different reasons. In the lines with increased B-type granule content, the early initiation of the B-type granules could allow more time for B-type granules to grow during grain development, resulting in larger B-type granules. However, in K2485, we observed that there was a significant reduction in the number of B-type granules. Thus, there are fewer granules competing for substates for granule growth, allowing each individual B-type granule to grow larger.

How the amino acid substitutions in MRC affect its function in granule initiation remains to be investigated. In the lines with increased B-type granule content (K2937, K4060, K0598, K2408 and K3889), the mutations (A178T, R255K, L289F, S503N and D581N, respectively) must have partially inhibited MRC function, since their granule size distributions shifted in the same direction as *mrc-1* with the premature stop codon mutation. Conversely, since the K2485 mutation had the opposite phenotype to *mrc-1*, the mutation (A625T) likely enhanced MRC function. The exact mechanism by which MRC acts is currently not known. However, we previously showed that MRC interacts directly with BGC1 (Chen et al. 2024). It is possible that the mutations in MRC specifically affect its ability to interact with BGC1, or with other granule initiation proteins. It is also possible that the mutations affect MRC protein stability. Protein destabilisation can inhibit function, whereas stabilising mutations can decrease the rate at which the protein is turned over, resulting in a longer protein lifetime. The latter is relevant since we proposed that MRC is expressed only during early grain development when it represses B-type granule initiation, and that the protein is not present during the later stages of grain development when B-type granules are initiating (Chen et al. 2024; Chen et al. 2023). A stabilising mutation may cause MRC protein to be retained during these later stages of grain development, thus inhibiting B-type granule initiation.

Although some significant changes in starch content and grain weight were observed among the lines (Supplemental Figure 2), the lines with the strongest alterations in B-type granule content (K2495, K0598 and *mrc-1*) had no change in these parameters. This suggests that the changes in granule size distributions can be achieved in the absence of changes in starch content.

### Granule size distributions affect starch physicochemical properties

Our RVA analyses demonstrated that the changes in granule size distribution induced by *Ta*MRC-6A mutations affect starch pasting properties, particularly peak viscosity and pasting temperature. Although the mutations affected B-type granule content and the size of A- and B-type starch granules, we determined that the effect on A-type granule size is likely to be the major contributor to the altered pasting properties. There was a strong positive correlation between peak viscosity and A-type granule size (Figure 3), which was still observed after removing the majority of B-type granules (Figure 4). Peak viscosity is one of the most important RVA parameters as it reflects the texture and thickness during starch cooking or gelatinisation. Our findings are consistent with the general trend that larger granules have higher viscosity (Li et al. 2021; Lindeboom et al. 2004). Also, given that A-type granules constitute the majority of the starch by volume in most of the lines examined, it is reasonable to expect that the pasting parameters are more influenced by the characteristics of the A-type granules. In addition to peak viscosity, A-type granule size (and thus B-type granule content) also strongly correlated with breakdown viscosity, peak time and pasting temperature (Supplemental Figure 4). All these parameters correlated poorly with B-type granule size. Rather B-type granule size correlated strongly with the setback viscosity, and to some degree with final viscosity.

Our study explored how pasting properties are affected by extensive, induced variation in granule size distributions within a single wheat cultivar. This contrasts the many previous studies that have investigated pasting properties in a narrower range of granule size variation existing between wheat cultivars, achieved through flour reconstitution, or following environmental treatments. Among these previous works, there is conflicting data on the effect of B-type granule content on pasting properties. Consistent with our study, higher B-type granule content in durum wheat was associated with lower peak viscosity, when variation in B-type granule content was achieved through crossing with *Triticum dicoccoides* or through reconstitution (Sissons and Egan 2023; Soh et al. 2006). This is also consistent with results in bread wheat: Wootton et al. (1998) found that B-type granule content negatively correlated with peak viscosity in a panel of Australian wheats, while Singh et al. (2008) also observed a similar negative correlation among different cultivars following water stress treatments. However, the opposite was found by Peterson and Fulcher (2001) and Dai Dai et al. (2009) – both genotypic variation in B-type granule content among wheat cultivars and variation induced by different watering treatments were positively correlated with peak viscosity. As discussed by Dai et al. (2009), it is likely that these contrasting results are due to pasting parameters being influenced by many other parameters, such as amylose and lipid content, which could also vary among genotypes and after environmental treatments. Indeed, Wootton et al. (1998) attributed the observed correlation between B-type granule content and peak viscosity to variation in amylose content, which also correlated with B-type granule content in the cultivars. An advantage of our study is that the variation in granule size distributions was induced by targeting a gene specifically involved in B-type granule initiation and is unlikely to alter other aspects of grain composition (Chen et al. 2024).

It should also be noted that the extent of variation in A-type granule size in these previous studies, and the degree to which that correlated with pasting properties is not known. The decreasing A-type granule size with increasing B-type granule content is a characteristic of *TtMRC-A1* mutations, and it cannot be assumed that A-type granule size correlates negatively with B-type granule content in a broader range of cultivars. Also, although our results suggest that A-type granule size is a stronger determinant of pasting properties than B-type granule content, we emphasise that this is only under the pasting conditions tested. The relative viscosities of fractionated A- and B-type granules are dependent on the concentration of starch used in the RVA, where A-type granules have higher viscosity than B-type granules at high solids content, while B-type granules have higher viscosity than A-type granules at lower solids content (Watanabe et al. 2024). It is therefore possible that we will see stronger contributions of B-type granule size and content to pasting parameters under different starch concentrations and pasting conditions.

The RVA parameters provide proof-of-concept that the altered granule size distributions achieved through *TtMRC-A1* affect starch physicochemical properties. The limitation of such laboratory tests is that they cannot always predict how the starches behave during industrial processing of food and other products. There can also be other benefits of having variation in granule size that are not directly linked to pasting parameters. For example, having lower B-type granule content in malt is beneficial for brewing (Bathgate and Palmer 1973). The digestibility of the starch may also be altered in the *mrc* mutants since A-type granules are digested more slowly in vitro than B-type granules (Colussi et al. 2021; Moon and Kweon 2024). Our set of *mrc* mutants provide an ideal panel that can be tested for improved quality in various applications of durum wheat, including pasta making.

### Applicability to other Triticeae crops

Given that *MRC* has orthologs in other Triticeae species (Chen et al. 2024), it is likely that it can also be targeted to alter starch granule size distributions in these species. However, it is an ideal gene target in durum wheat because the B-genome homeolog became a pseudogene shortly after the hybridisation that gave rise to tetraploid wheat (Chen et al. 2024), allowing changes in granule size to be induced by altering the single *TtMRC-A1* locus. We expect that a similar benefit will be seen in barley, a diploid Triticeae that also has only one gene encoding MRC. However, in hexaploid bread wheat, there are two active homeologs of *MRC* on chromosomes 6A and 6D, and it is likely both homeologs need to be targeted to induce substantial changes to the granule size distribution. Missense mutations may be induced using chemical mutagenesis and identified using TILLING approaches, or induced using CRISPR-based base editing. The mutations that significantly increased (A178T, R255K, L289F, S503N and D581N) and decreased B-type granule content (A625T) were all in amino acids that are conserved in all wheat and barley MRC sequences examined (Supplemental Figure 5), suggesting that the same mutations can be targeted in bread wheat and barley. Interestingly, the missense mutations that led to the strongest increase (L298F) and decrease (A625T) in B-type granule content were in amino acids that were also conserved in the Arabidopsis and rice proteins, suggesting that these amino acids may play an important role in MRC function in a broader range of species.

## Acknowledgements

We thank Dr. Rose McNelly (John Innes Centre) for assistance in preparing figures. We also thank JIC Horticultural Services for providing growth facilities and maintenance of plant material, JIC Germplasm Resource Unit for providing access to the wheat TILLING mutants, JIC Bioimaging for providing access to microscopes, and the JIC Genotyping service for conducting KASP genotyping.

## Statements & Declarations

### Author contributions

DS conceived the study. BF and DS designed the research. BF and JC performed the research. All authors analysed data. DS wrote the article with input from all authors.

### Data availability

The data underlying this article are available in the article and in its supplementary material.

## Supplemental Figures

**Supplemental Figure 1:**
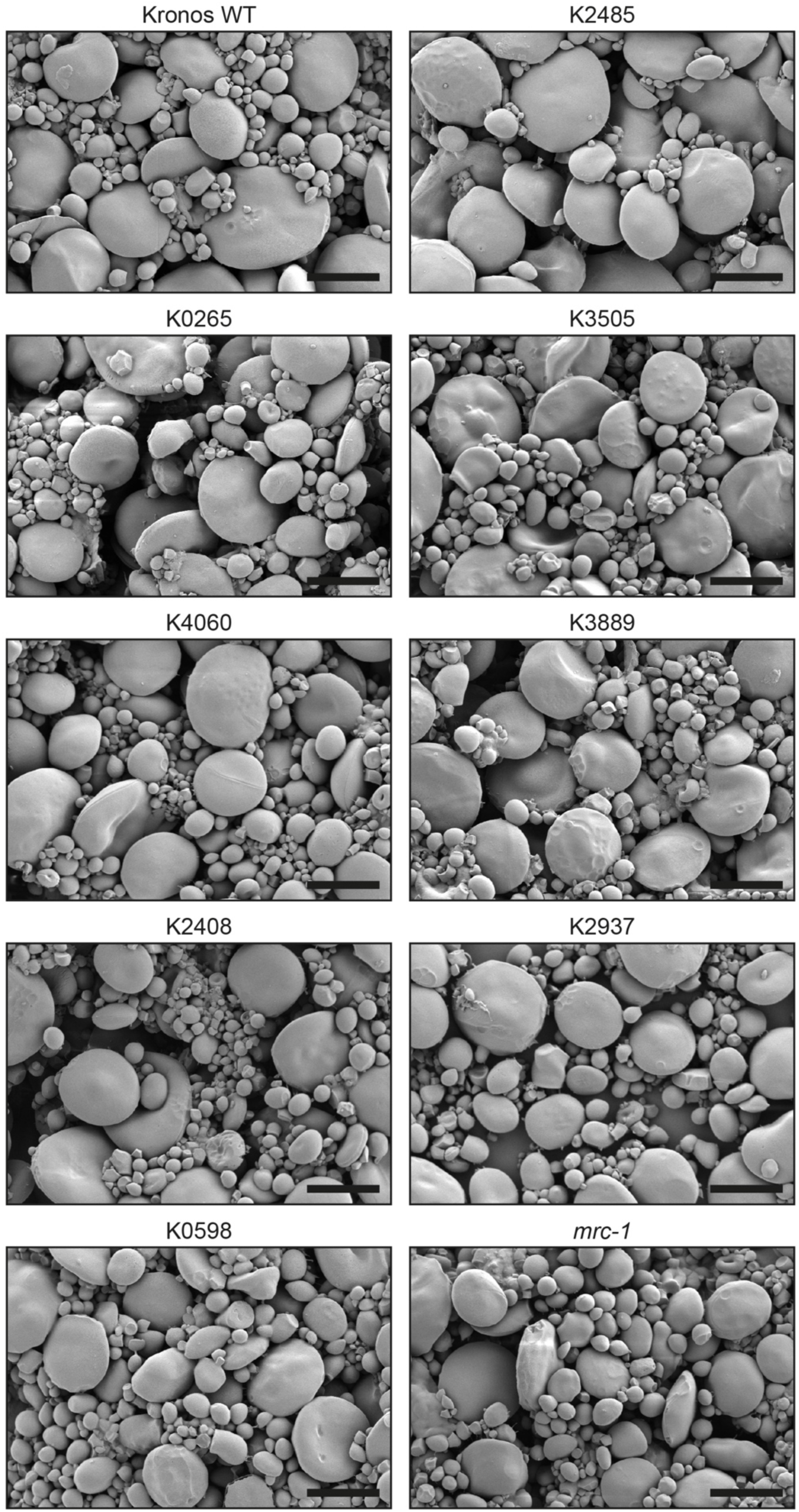
Starch granule morphology of Kronos TILLING mutants in Experiment 2. Purified starch granules were imaged with a Scanning Electron Microscope. Bars = 20 µm.

**Supplemental Figure 2:**
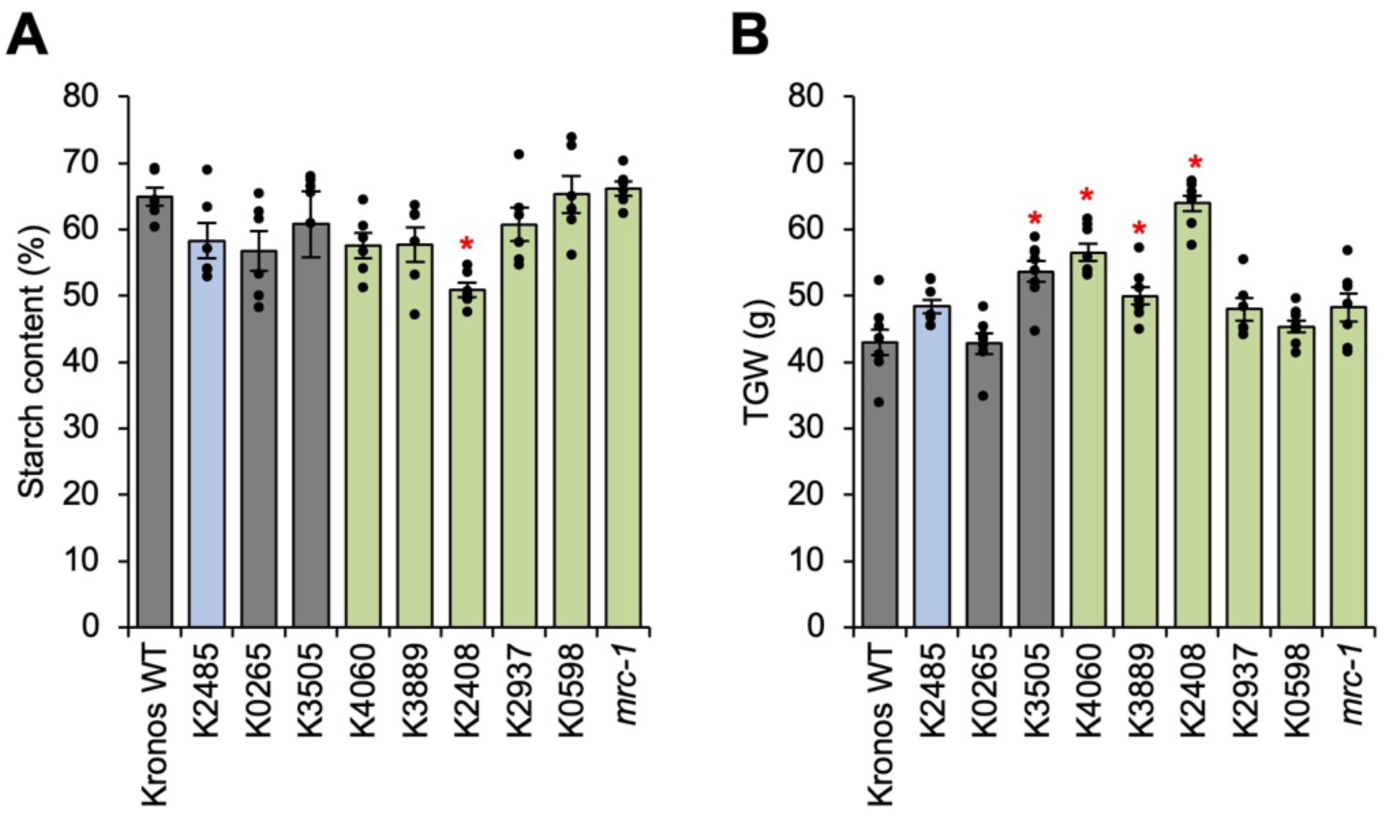
Starch content and grain weight of mutants examined in Experiment 2. (A) Total starch content was determined by enzymatic quantification (B) Thousand Grain Weight (TGW) was determined using the Marvin seed analyser. Values are the mean ± standard error of the mean (SEM) from *n*=6 (panel A) or *n*=6-8 (panel B) biological replicates, where each replicate (shown as an individual data point) was prepared from grains harvested from a separate plant. Values marked with an asterisk are significantly different to the Kronos wild type (WT) under a one-way ANOVA and Tukey’s HSD test (p < 0.05).

**Supplemental Figure 3:**
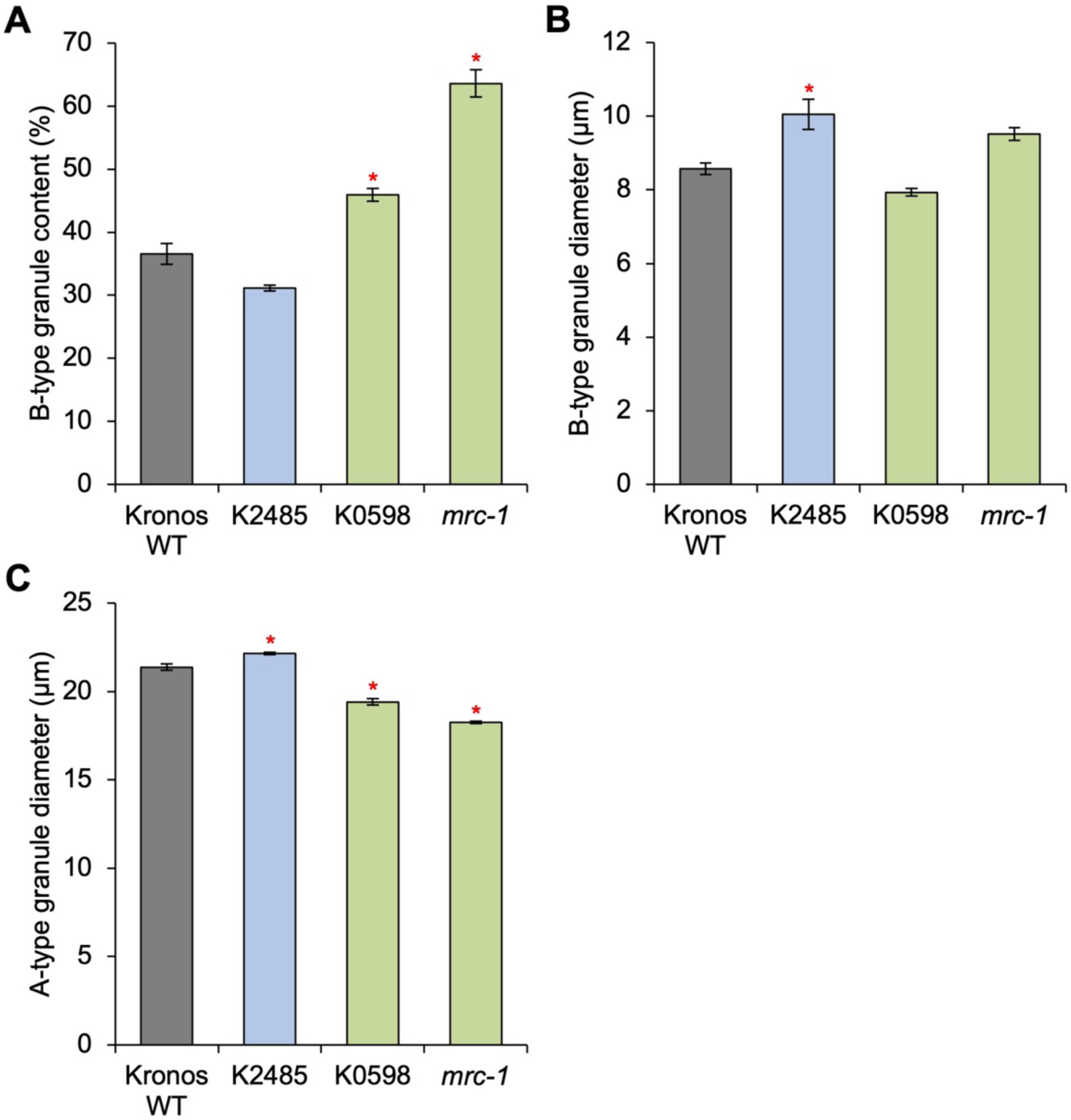
Granule size parameters on large-scale starch extracts used for the Rapid Visco Analysis. Coulter Counter analyses were carried out on starches (shown in Figure 3A), and granule size parameters were calculated using curve-fitting analyses. (A) B-type granule content (by relative volume). (B) Mean B-type granule diameter. (C) Mean A-type granule diameter. Values are the mean ± standard error of the mean (SEM) from *n* = 3 replicate measurements, and those marked with an asterisk are significantly different to the Kronos wild type (WT) under a one-way ANOVA and Tukey’s HSD test (p < 0.05).

**Supplemental Figure 4:**
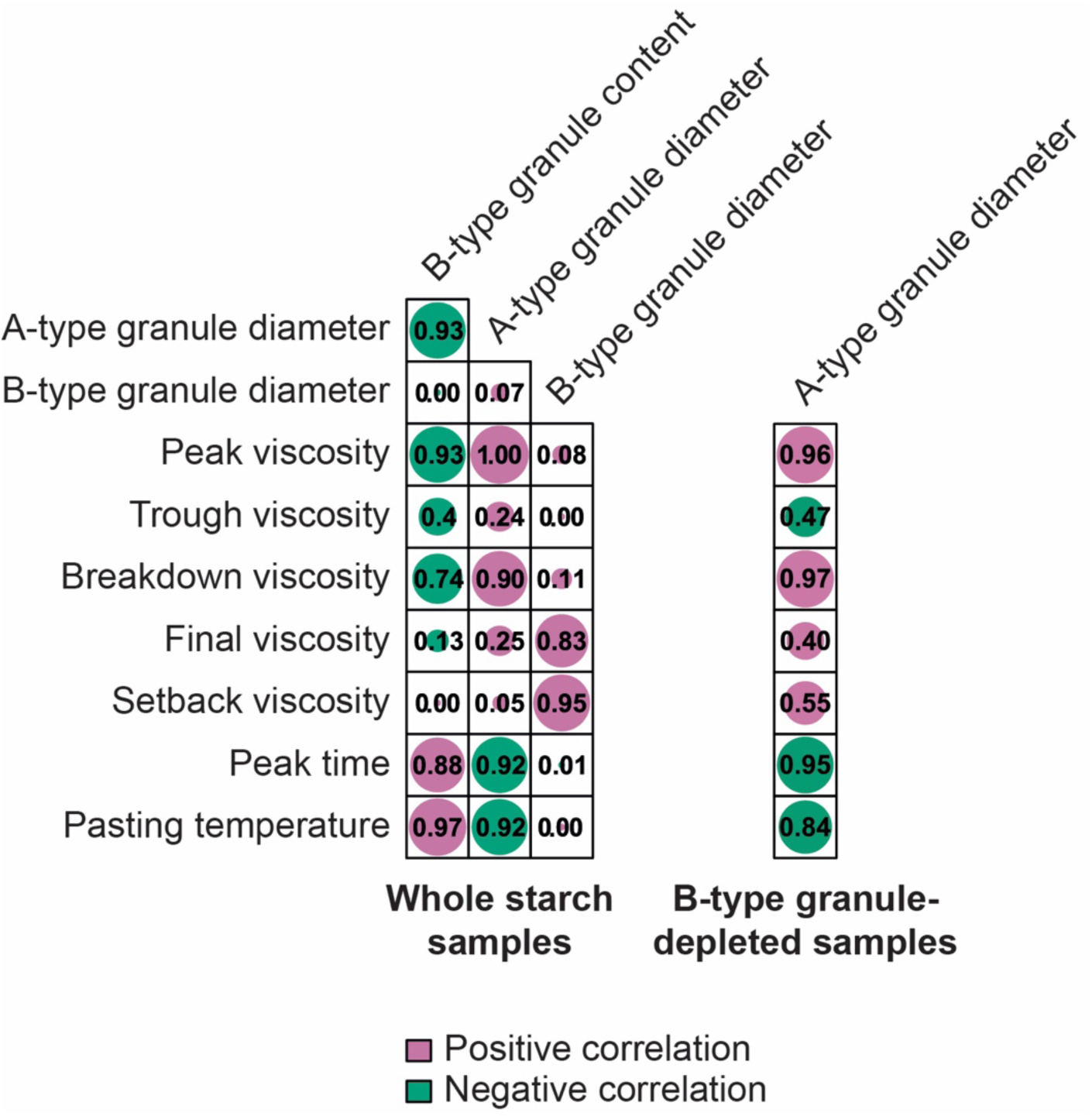
Plot of R^2^ values between pairwise linear regressions for granule size and pasting parameters. Numbers represent the R^2^ value, while the size of the dot is proportional to the value. Positive correlations are marked with pink dots, while negative correlations are marked with green dots.

**Supplemental Figure 5:**
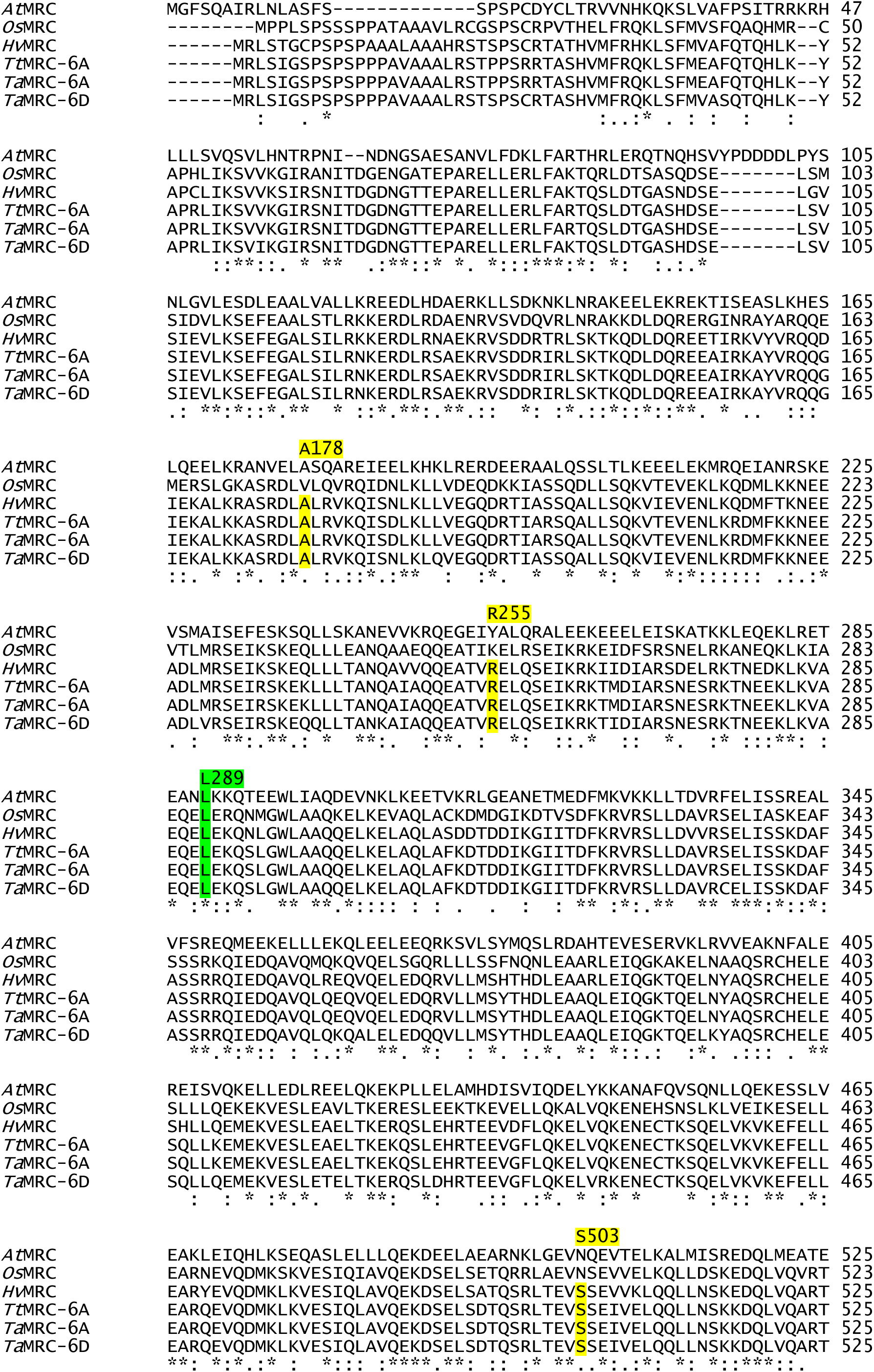
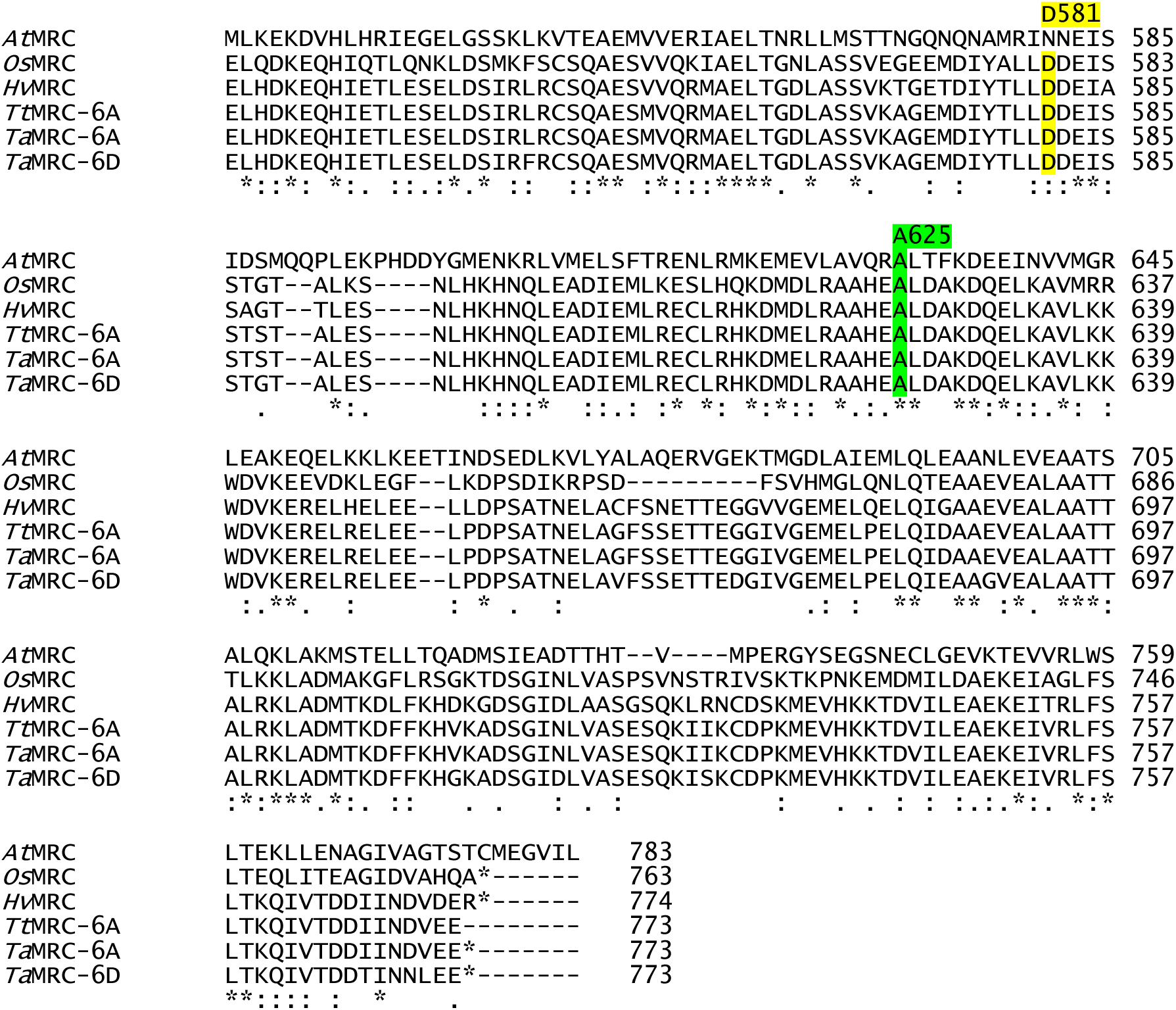
Amino acid alignment of wheat and barley MRC proteins. The alignment was conducted using Clustal Omega including MRC sequences from durum wheat (*Tt*MRC-6A, TRITD6Av1G081580.1), bread wheat (*Ta*MRC-6A, TraesCS6A02G180500.1; *Ta*MRC-6D, TraesCS6D02G164600.1), barley (*Hv*MRC, HORVU6Hr1G036020.1), rice (*Os*MRC, LOC_Os02g09340.1) and Arabidopsis (*At*MRC, At4g32190). The positions of the amino acids where mutation increased B-type granule content (A178T, R255K, L289F, S503N and D581N), and the one where mutation decreased B-type granule content (A625T) are highlighted. Green highlight indicates amino acids conserved in all sequences examined, while yellow highlight indicates amino acids that are conserved in all wheat and barley sequences.

**SUPPLEMENTAL TABLE 1.**
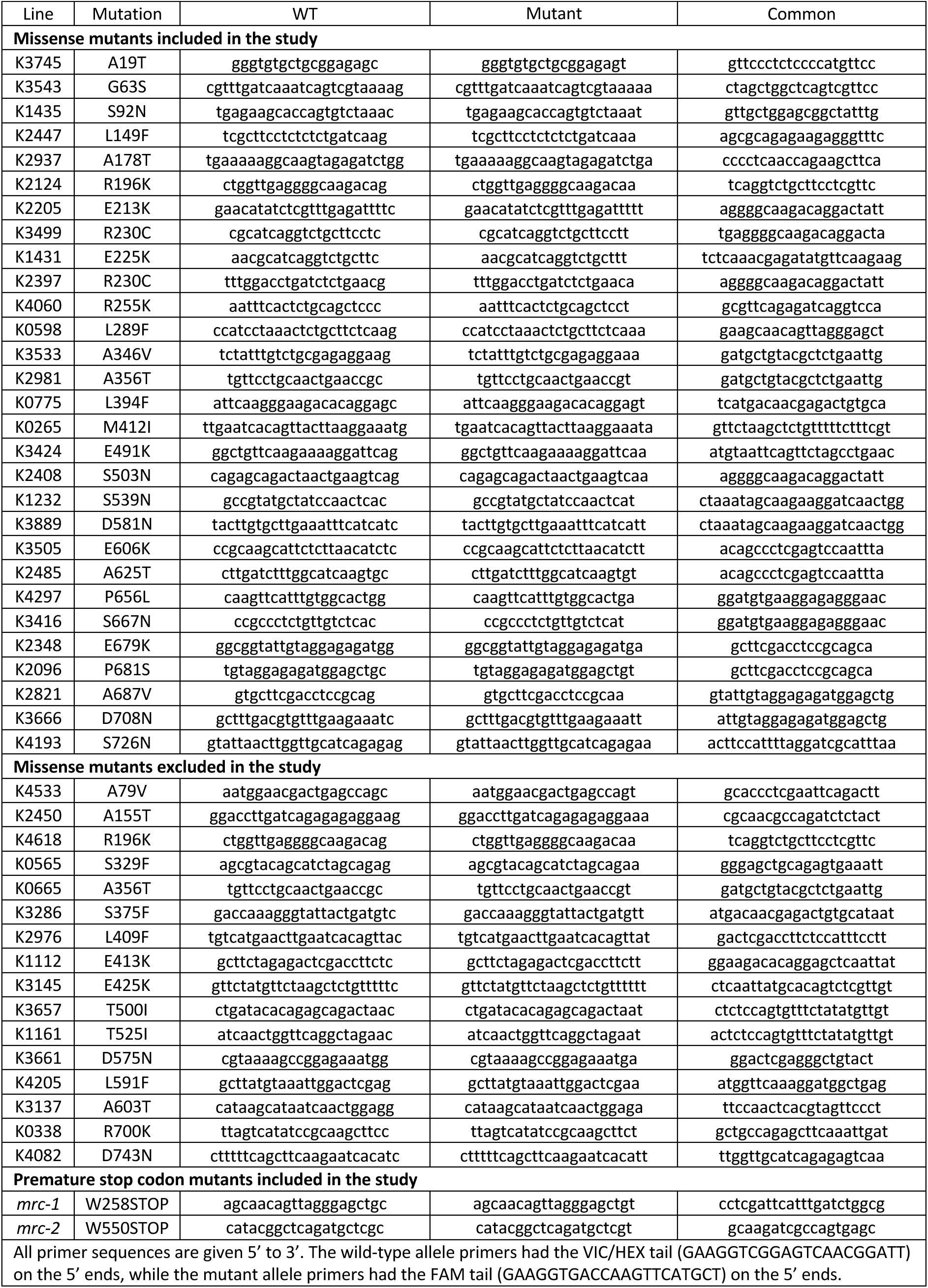
KASP markers used in this study.

